# Molecular characterization of a flatworm Girardia isolate from Guanajuato, Mexico

**DOI:** 10.1101/2020.07.01.183442

**Authors:** Elizabeth M. Duncan, Stephanie H. Nowotarski, Carlos Guerrero-Hernández, Eric J. Ross, Julia A. D’Orazio, Clubes de Ciencia México Workshop for Developmental Biology, Sean McKinney, Mark C. McHargue, Longhua Guo, Melainia McClain, Alejandro Sánchez Alvarado

**Author notes:** Equal contributors (Elizabeth M. Duncan, Stephanie H. Nowotarski).

## Abstract

Planarian flatworms are best known for their impressive regenerative capacity, yet this trait varies across species. In addition, planarians have other features that share morphology and function with the tissues of many other animals, including an outer mucociliary epithelium that drives planarian locomotion and is very similar to the epithelial linings of the human lung and oviduct. Planarians occupy a broad range of ecological habitats and are known to be sensitive to changes in their environment. Yet, despite their potential to provide valuable insight to many different fields, very few planarian species have been developed as laboratory models for mechanism-based research.

Here we describe a previously undocumented planarian isolate, *Girardia sp*. (Guanajuato). After collecting this isolate from a freshwater habitat in central Mexico, we characterized it at the morphological, cellular, and molecular level. We show that *Girardia sp*. (Guanajuato) shares features with animals in the *Girardia* genus but also possesses traits that appear unique to this isolate. By thoroughly characterizing this new planarian isolate, our work facilitates future comparisons to other flatworms and further molecular dissection of the unique and physiologically-relevant traits observed in this *Girardia sp*. (Guanajuato) isolate.

## BACKGROUND

Biologists have been fascinated by planarian flatworms for centuries, in large part because of their remarkable ability to regenerate missing tissues and body parts [1]. Hundreds of diverse planarian species have been identified in numerous locations across the globe and in various environments, including freshwater [2], marine [3], and terrestrial habitats [4]. Despite their morphological and ecological diversity, most planarians have at least some regenerative capacity, although there is a wide range of species-specific abilities. At one extreme, some freshwater planarians can recreate an entirely new animal from a small fragment of amputated tissue [5]; at the other end of the spectrum, many marine species exhibit very limited regeneration *e.g*., only within a restricted anatomical region [6,7]. This variation in regenerative abilities and ecological habitats not only makes planarians an excellent model for studying the cellular mechanisms underlying animal regeneration, but also provides an opportunity to assess the impact of various environmental factors on this complex developmental and reproductive process. For example, excess amounts of industrial pollutants such as ferric iron and organophosphorus pesticides have been shown to affect functional head regeneration [8–10]. Such studies provide valuable insight into how changes in the environment, including increased exposure to specific chemicals, can impact both the immediate health of local animals and their ability to maintain their population.

Although their impressive regenerative capacity sets them apart, planarians also bear several biological features that are morphologically and functionally analogous to those of other animals, including humans. One important such feature is the planarian epidermis, which comprises a monostratified mucociliary epithelium that is highly similar to the epithelia of human lungs and oviducts [11]. Planarians also possess primitive nephron units, or protonephridia, which share striking structural, functional, and transcriptional identity with their counterparts in human kidneys [12]. In short, planarians are a powerful, tractable, *in vivo* research organism for studying many fundamental cellular structures, functions, and processes.

Given the diversity of planarian species and their differences in regenerative capacity, morphological features, and interactions with the local environments, comparative studies offer enormous potential for uncovering the genomic and cellular mechanisms underlying these unique characteristics. However, only a few planarian species, mainly *Schmidtea mediterranea* (*S.med*) and *Dugesia japonica*, have been sufficiently developed into models with established tools and infrastructure for such analyses. In order to expand the power of comparative approaches and the number of distinct biological features that can be compared and dissected, we need to identify and thoroughly characterize new planarian species and isolates.

Here we describe a new isolate that we name *Girardia sp*.(Guanajuato), abbreviated *G.sp*.(Guanajuato). We characterize its basic anatomy and capacity for regeneration and describe features of its outer epithelium and stem cell population that distinguish *G.sp*.(Guanajuato) from other species and isolates. We also assembled a *G.sp*.(Guanajuato) transcriptome, which will greatly facilitate gene expression analyses and forward genetic experiments in future studies. This thorough characterization of *G.sp*.(Guanajuato) will both stimulate further investigations into its intriguing features and encourage the discovery of more planarian species and isolates.

## RESULTS

### Animal Collection

Undergraduate students of a Developmental Biology Workshop organized by Clubes de Ciencia (CdeC) (https://www.clubesdeciencia.mx/) hosted at the University of Guanajuato in Guanajuato, México, embarked on a field expedition to find and study native planarians. We visited the Parque Bicentenario in the southeastern part of Guanajuato City, Mexico to prospect for animals in August of 2016 (Figure 1A, red arrowhead). The Parque Bicentenario is home to the Presa San Renovato (Saint Renovato Dam), which controls the flow of several local springs and small rivers. We found the new isolate in a shallow pool of freshwater with minimal flow opposite the dam (Figure 1B). We observed an aggregation of approximately 300 animals in this pool (Figure 1C) and, upon closer inspection, noted that the majority of animals appeared to be around 1 centimeter in length with dark pigmentation and locomoting in an undulating manner (Figure 1D). We collected approximately 60 animals and transported them back to the laboratory for further characterization (Figures 2-7).

**Figure 1.**
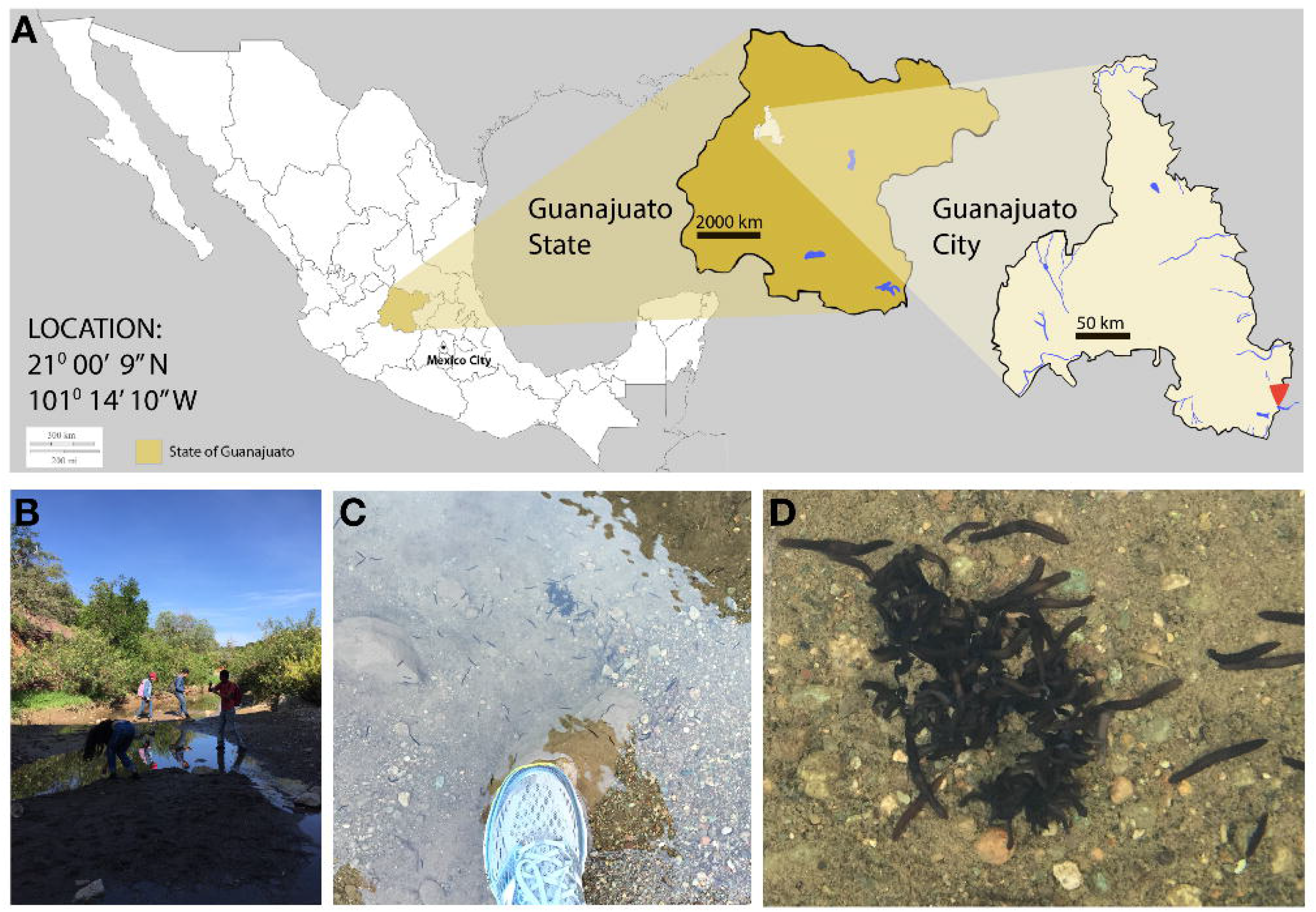
A previously uncharacterized planarian isolate was collected in Mexico. (A) A map showing the location of the Presa San Renovato (red arrowhead), a dam within the Parque Bicentenario where the newly named species of planarians was collected. Magnified images show the site is located at the southeastern edge of Guanajuato City in the state of Guanajuato, Mexico. (B-D) The collected isolate *G.sp*.(Guanajuato), was found at the bottom of a shallow pool of freshwater near the dam.

**Figure 2.**
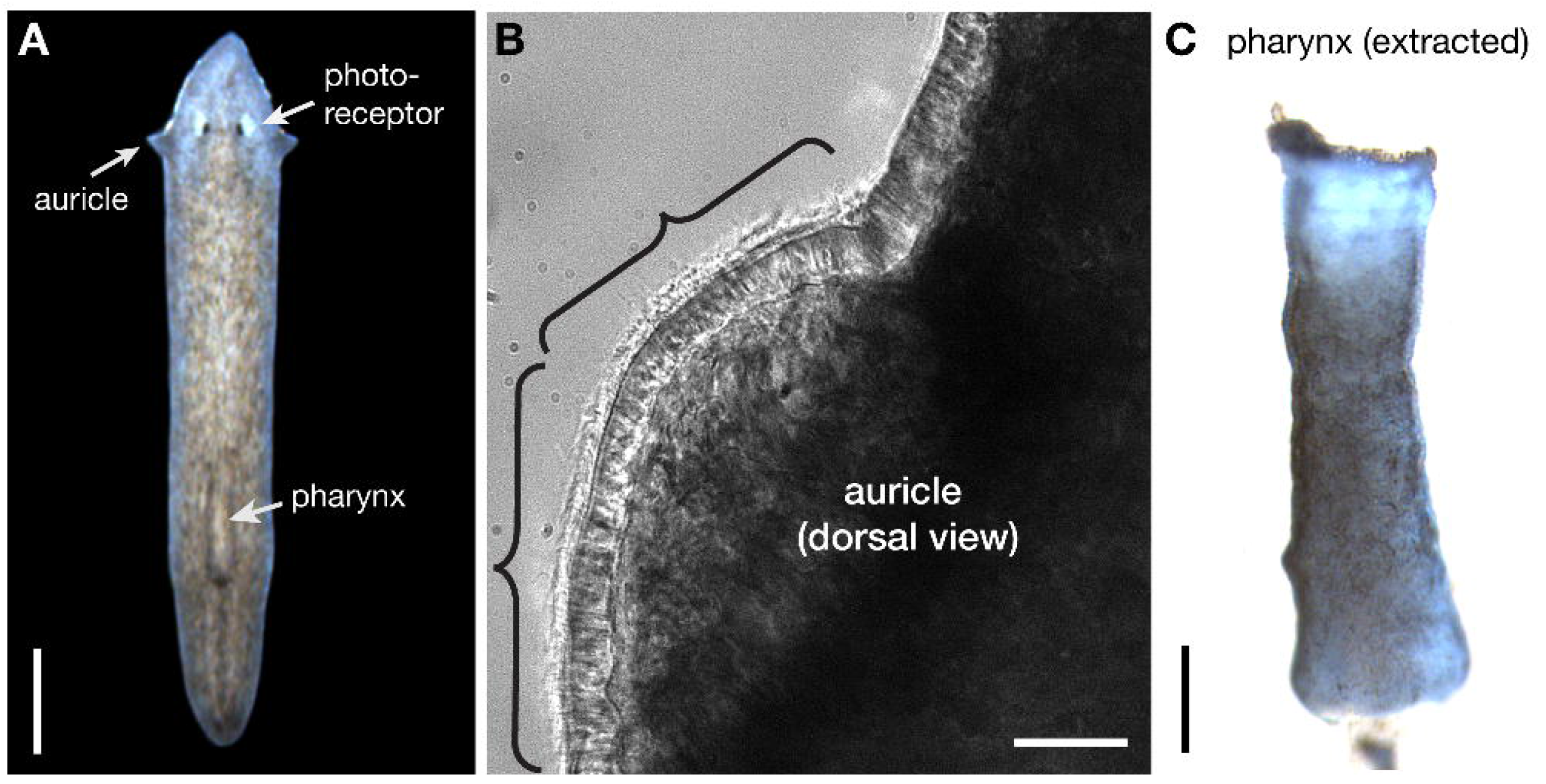
*G.sp*.(Guanajuato) exhibit standard morphological features associated with *Girardia* planarians. (A) Live image of a single *G.sp*.(Guanajuato) worm after transporting them back to the laboratory. As with other planarians in the *Girardia* genus, *G.sp*.(Guanajuato)has prominent, ciliated auricles (A,B) and a pigmented pharynx (C). Scale bars = 500μM in A, 20μM in B, 200μM in C. Brackets in (B) indicate cilia projecting from the epidermal cells that cover the auricle.

**Figure 3.**
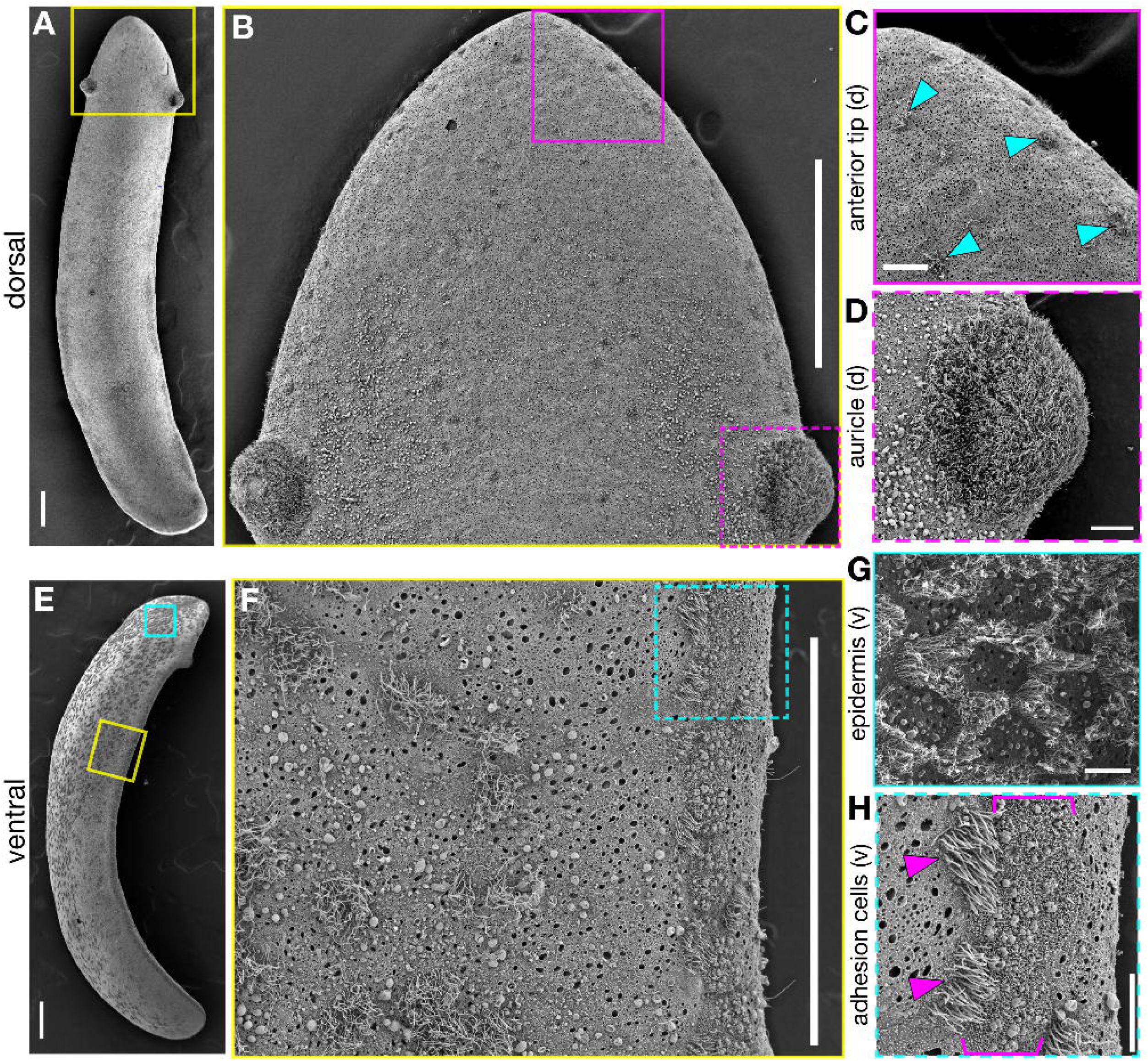
The epidermis of *G.sp*.(Guanajuato) planarians has a distinct pattern of ciliated cells. (A) A Scanning Electron Microscopy (SEM) image of a *G.sp*.(Guanajuato) planarian, viewed from the dorsal side. As seen in (A-D), the dorsal epidermis is largely non-ciliated (sparse individual ciliated cells are indicated by cyan arrowheads) except for the auricles, which are densely ciliated on the dorsal and lateral sides (B,D). The ventral epidermis (E-H) exhibits a distinct pattern of both ciliated cells and non-ciliated cells, which differs from the densely ciliated ventral epidermis seen in many other planarian species. As seen in (F,H), the decreased density of ciliated cells in the ventral epidermis allows a line of adhesion gland cells (brackets) to be observed easily by SEM. Individual ventral ciliated cells that lie along the adhesion glands are indicated by magenta arrowheads. Scale bars = 200μM in A,B,E,F; 20μM in C,D,H; 10μM in G.

**Figure 4.**
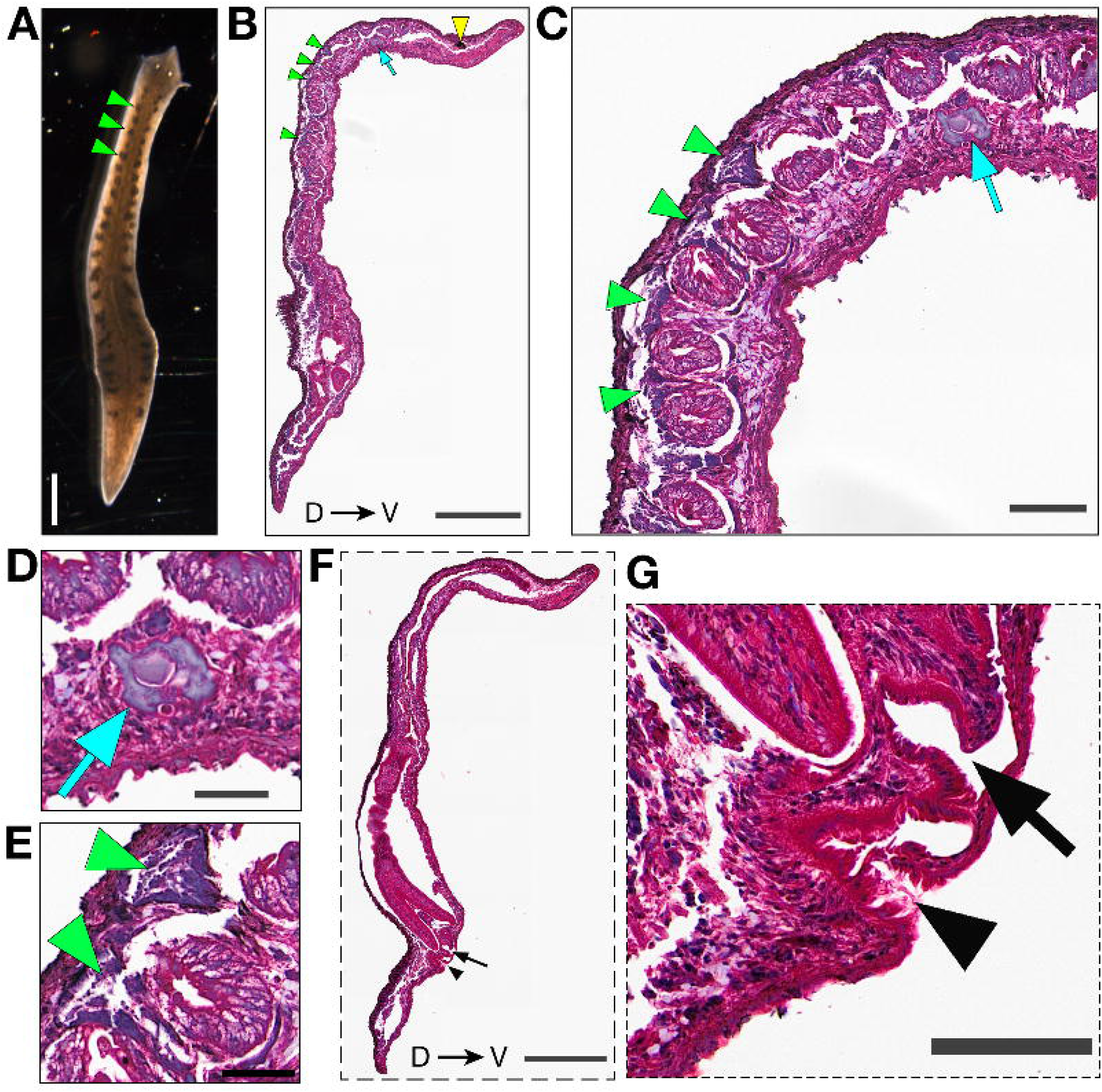
*G.sp*.(Guanajuato) planarians show sexualized morphology at very low frequency in the lab. (A) Darkfield image of a rare sexualized animal revealing a clear pattern of dark nodes (green arrowheads) visible dorsolaterally. (B,F) H+E stained histological sections representing two different sagittal planes of an animal with a sexualized appearance. One section (B) reveals an ovary (cyan arrow, B-D) on the ventral side of the worm posterior to the photoreceptors (yellow arrowhead); it also includes testes on the dorsal side (green arrowheads in B,C,E) nested between gut branches. A second more medial sagittal section (F,G) reveals a gonopore (black arrowhead) and atrium (black arrow). Scale bars = 1mm in A, 0.5mm in B and F, 100μm in C and G, 50μm in D and E.

**Figure 5.**
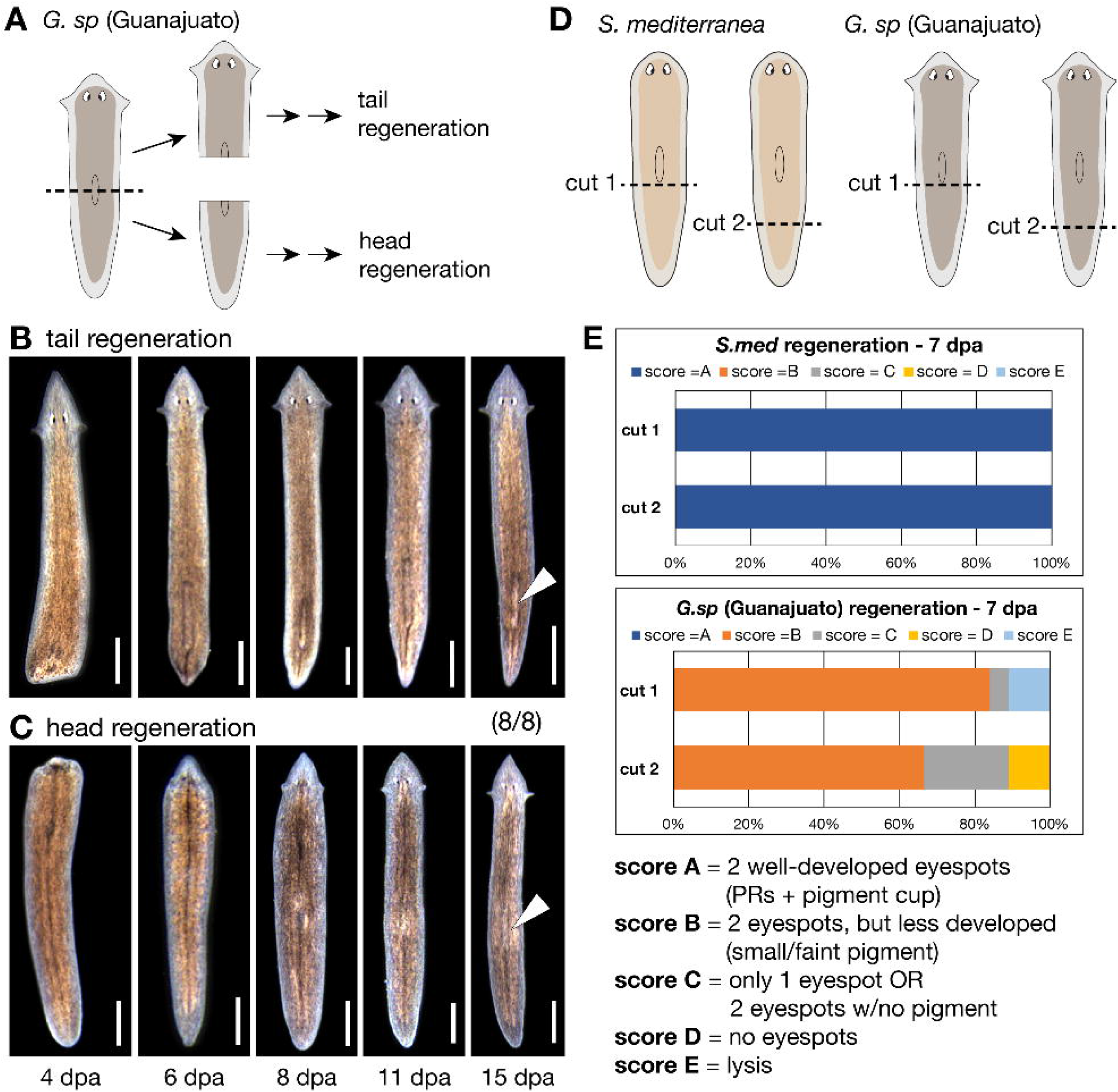
*G.sp*.(Guanajuato) have high regenerative capacity. (A) Schematic of the amputation strategy used to test the regeneration capacity of newly isolated worms. Amputated *G.sp*.(Guanajuato) fragments demonstrate a rapid and robust capacity for both new tail regeneration (B) and new head regeneration (C), including the formation of a new pharynx in both fragments (white arrowheads). (D) Schematic of the amputation strategy used to compare regeneration capacity in *G.sp*.(Guanajuato) versus the well-characterized species *Schmidtea mediterranea* (*S.med*). (E) Plot showing percentage of worms for each species with the indicated regeneration score 7 days post-amputation (dpa). Scale bars = 1mm in A-C, 100μm in F.

**Figure 6.**
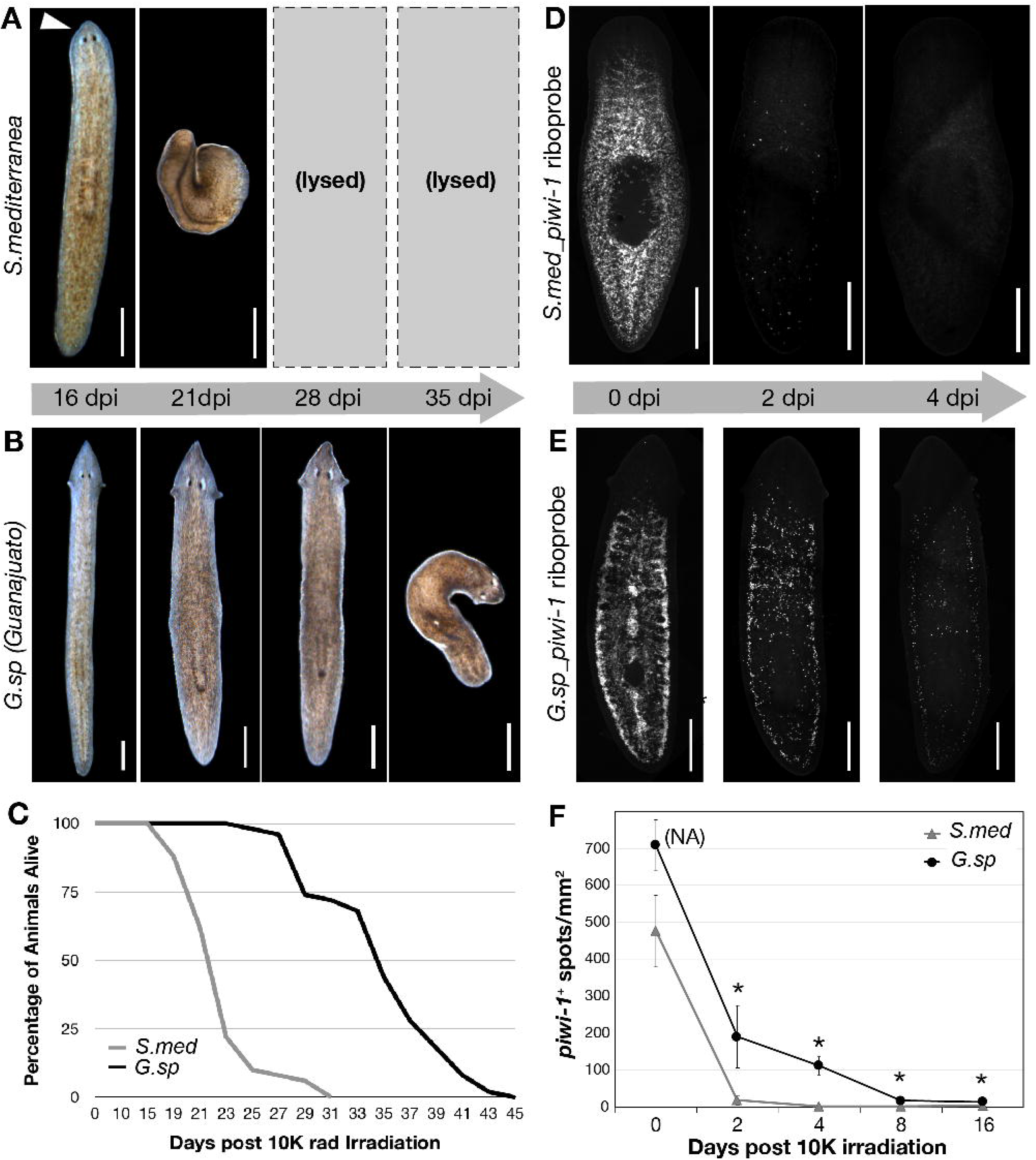
*G.sp*.(Guanajuato) planarians have a higher tolerance for □-radiation than *S.med* planarians. 50 worms per *S.med* (A) and *G.sp*.(Guanajuato)(B) were exposed to the same dose of □-radiation (10K rads) and monitored over time. Images are representative of the phenotype seen in most worms at a given time point (days post-irradiation; dpi). (C) Survival data from the experiment represented in A and B was recorded at the indicated days post radiation treatment and plotted as shown. (D) *in situ* hybridization images of representative *S.med* worms or (E) *G.sp*.(Guanajuato) worms fixed at the indicated time points post □-radiation and stained with specific *piwi-1* riboprobes. (F) Quantitation of *piwi-1^+^* cells for the experiment shown in D and E; n=4-5 worms at each time point. NA = density of *piwi-1* signal is too high at the indicated time point for accurate quantitation. * = p<0.02. All scale bars = 500μM.

**Figure 7.**
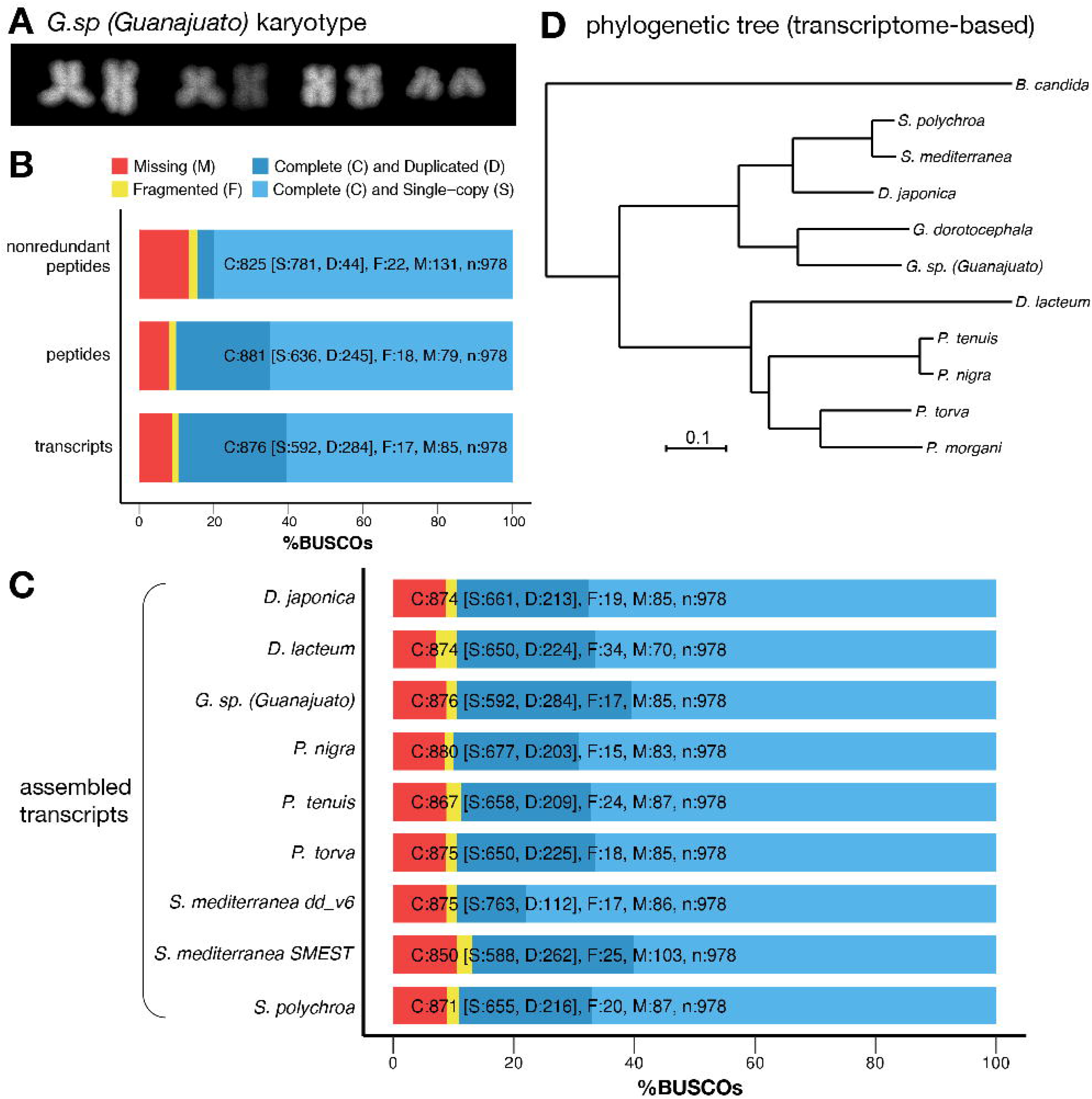
*G.sp*.(Guanajuato) are diploid planarians in the *Girardia* genus. (A) Karyotype of cells in *G.sp*.(Guanajuato) tail tissue. (B) Transcriptome completeness evaluation of the newly assembled *G.sp*.(Guanajuato) transcriptome using 978 metazoan BUSCOs. (C) Comparison of the newly assembled *G.sp*.(Guanajuato) transcriptome to 8 other assembled transcriptome downloaded from the PlanMine database[33]. (D) Phylogenetic tree created using 3,163 concatenated single-copy putative orthologs from 11 triclad flatworm species. Scale bar unit = expected number of substitutions per site.

### Morphological Characterization

In order to begin to assign this *G.sp*.(Guanajuato) planarian to a genus, we first characterized their external morphology at low resolution (Figure 2). We immediately noted the presence of two prominent eyespots (or photoreceptors), a triangular head with pointed auricles, and an oblong dorsal patch of lightly pigmented tissue that is indicative of an internal pharynx (Figure 2A). Upon inspection at higher magnification, we observed that the auricles are densely ciliated (Figure 2B, brackets) with highly motile cilia (Supplemental Movie S1). Several genera of freshwater planarians feature pronounced, ciliated auricles, including *Dugesia, Phagocata* and *Girardia* [13–15]. Yet the pointed shape of the *G.sp*.(Guanajuato) head and auricles largely exclude *Phagocata* as a possibility. To distinguish between *Dugesia* and *Girardia*, we presented animals with food and monitored as they extruded their pharynges to eat. In all worms, we observed a single, pigmented pharynx (Figure 2C). As all other characterized planarians with pigmented pharynges are classified as *Girardia* and this trait is considered apomorphic for the genus [16], this new isolate likely belongs to the *Girardia* genus. This assignment is also supported by the fact that only four genera are documented to be endemic to the Neotropical region: *Romankenkius, Opisthobursa, Rhodax* and *Girardia*, with *Girardia* species accounting for the majority [17].

Like other planarians of this genus, the pharynx is covered by an infranucleated and heavily ciliated epithelia that is directly superficial to longitudinal muscle fibers which are in turn superficial to circular muscle fibers (Supplemental Figure 1B). The internal muscle layers are repeated with an inner longitudinal muscle layer and an outer and well-developed circular muscle fiber layer (Supplemental Figure 1C). In addition to the pharynx muscle layers, we noted that the body wall muscle layers consisted of thin circular fibers just underneath the basal lamina, followed by thin longitudinal fibers underneath, then diagonal fibers, and finally (at the most internal) thick longitudinal fibers that appear to be more developed ventrally (Supplemental Figure 1 D,E).

As many planarian characterizations also note the length of the pharynx in relation to the body length, we also measured this ratio. After imaging eight live, individual animals (as in Figure 2A), surgically removing their pharynges, and imaging the corresponding pharynges (as in Figure 2C), we measured all body and pharynx lengths and calculated their ratios (Supplemental Table 1). Our measurements show that *G.sp*.(Guanajuato) pharynges are approximately 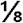 of body length, which is both consistent across animals of different body lengths and significantly smaller than what is reported for other *Girardia* (¼ - □ body length; Supplemental table 2).

Planarian locomotion is driven by the concerted beating of motile cilia that protrude from cells in the ventral epidermis [14,18]. When *G.sp*.(Guanajuato) migrate towards food in a laboratory dish, they frequently perform a peristaltic-like motion (Supplemental Movie S2) rather than the canonical smooth glide associated with other common laboratory species. This motion was also noted during collection in the field. As this motion is reminiscent of the inch-worming phenotype exhibited by *S.med* animals after RNAi depletion of essential cilia proteins [19,20], we decided to examine the outer mucociliary epidermis of these animals. To characterize all potentially important exterior features of this tissue, we used scanning electron microscopy (SEM) to capture images with detailed resolution. SEM revealed that the dorsal epidermis of *G.sp*.(Guanajuato) has two distinct regions: 1) the auricles, which are covered by densely ciliated cells (Figure 3A, B, D); and 2) the remaining dorsal surface, which is non-ciliated except for single, sparsely distributed, ciliated cells that appear in lateral stripes (Figure 3B, C). Examination of the ventral surface by SEM (Figure 3E-H) revealed clustered patches of multiciliated cells that span the entire ventral epidermis, although with higher density at the anterior end. In some regions of the animal these patches are organized in a distinct checkered pattern of single ciliated and non-ciliated cells (Figure 3G; Supplemental Figure 2). Although this unique pattern of ventral cilia does correlate with and possibly explains the motility differences observed for *G.sp*.(Guanajuato) animals, we were still surprised by this finding. Most other planarian species, including *S.med* and *Girardia tigrina* [11,14], maintain a dense and apparently uniform distribution of multiciliated cells across their ventral epithelia such that it resembles a lawn of cilia rather than patches.

Our SEM images indicated this pattern was created by the specification of distinct cell types rather than the random loss of cilia due to mechanical stress or osmotic shock. To validate this hypothesis, we also examined the epithelial cells of a transverse cross-section through the auricles at subcellular resolution (Supplemental Figure 3). We readily observed a clear distinction between ciliated cells and non-ciliated cells, most notably because the non-ciliated cells contain rod-like rhabdite structure in both the dorsal and ventral epidermis whereas these structures are largely absent in ciliated cells. This clear separation between rhabdite-enriched, non-ciliated cells and densely ciliated cells (lacking rhabdites) is observed across the epidermis (Supplemental Figure 3), supporting the idea that the patchy pattern of ciliated cells we describe in *G.sp*.(Guanajuato) is likely a reflection of differences in cell fate rather than the loss of cilia during animal handling or processing.

While maintaining *G.sp*.(Guanajuato) in the lab, we observed multiple instances of reproduction by fission, which is well documented in the asexual *S.med* biotype [21] but rarely seen in the sexual strain of *S.med* or other sexual planarian species. We were therefore surprised to note that a small number of animals in the *G.sp*.(Guanajuato) colony had two ventral openings, suggesting these animals were sexualized. We also observed putative testes when examining these rare worms using darkfield microscopy (Figure 4A). We detected these putative sexual animals at an extremely low frequency: n=4 animals in the fall of 2017 and n=10 animals in the fall of 2018 with none documented in the following three years. After sequestering these putative sexual animals together for extended periods of time, we never observed any copulatory behavior or eggs, suggesting at best an immature state of sexualization. To determine if these animals indeed possessed sexual organs, we submitted animals from our 2018 collection for histological analysis. *G.sp*.(Guanajuato) animals were sectioned sagittally in order to obtain lateral planes containing both an ovary (Figure 4B-D cyan arrows) and testis (Figure 4B,C,E green arrowheads). As seen in the sexual biotype of *S.med*, we observed ovaries on the ventral side of *G.sp*.(Guanajuato) sections in a position posterior to the photoreceptors and just behind the cephalic ganglia (Figure 4B,C, cyan arrow). Sagittal sectioning also revealed details of the medial copulatory region, including a gonopore (Figure 4F,G black arrowhead) and what appears to be an immature copulatory apparatus with an infranucleated epithelial lining and lumen of an atrium (Figure 4F,G black arrow and Supplemental Movie S3). These data suggest that *G.sp*.(Guanajuato) animals may be capable of transitioning from a fissiparous (asexual) state to a sexualized state, although it is difficult to conclude without evidence of functional sexual reproduction.

### Regenerative Capacity

Many planarians have the capacity to regenerate lost tissues, although this ability varies across species and isolates. Planarians with limited regenerative abilities often cannot regenerate new heads if amputated near or below the pharynx [22–24]. We first assayed the regenerative capacity of *G.sp*.(Guanajuato) animals by bisecting individual animals transversely through the pharynx and monitoring their recovery over the course of 15 days (Figure 5A-C). All head and tail *G.sp*.(Guanajuato) fragments demonstrated successful wound healing and survived throughout the course of the experiment (8/8 animals). At four days post amputation (dpa), we observed early signs of tissue regeneration in both head (Figure 5B) and tail (Figure 5C) fragments, including the formation of a small, non-pigmented blastema in tail fragments (Figure 5C). After six days, tail fragments showed signs of newly regenerated photoreceptors (e.g., two spots of dark pigment at the anterior wound site) and after 8 days the eyespots were clearly and completely formed (Figure 5C). All worm fragments had fully regenerated within fifteen days. These data clearly show that *G.sp*.(Guanajuato) planarians have a robust regenerative capacity when cut through the pharynx.

To assess the regenerative potential of *G.sp*.(Guanajuato) further, we then performed amputations in both the highly regenerative *S.med* planarian species and *G.sp*.(Guanajuato) at two amputation planes below the pharynx (Figure 5D). This experiment revealed that although *G.sp*.(Guanajuato) animals are capable of regeneration when injured below the pharynx, their capacity is significantly less robust as compared to *S.med* (Figure 5E and Supplemental Figure 4). *G.sp*.(Guanajuato) fragments not only regenerated more slowly than *S.med* fragments overall, but also showed more variability in regenerative success (Figure 5E), results that are similar to those reported for another *Girardia* species [25]. Our experiments did not indicate that animal body length is the major factor driving *G.sp*.(Guanajuato) regeneration success, although this factor will be evaluated more thoroughly in future studies. We did, however, observe that *G. sp* (Guanajuato) regenerating fragments may be more sensitive to water conditions than *S.med:* in some experiments when both species of animals were maintained and amputated in 0.5g/L Instant Ocean (versus 1X Montjüic planarian water), regenerating *G.sp*.(Guanajuato) fragments had higher rates of lysis (data not shown) whereas *S.med* regeneration was consistently successful in either water condition.

### Stem cell population

The regenerative capacity of planarians depends on a population of adult stem cells, also known as neoblasts, that bear many unique properties [26]. Not only do these stem cells self-renew and differentiate as needed during both regeneration and normal homeostasis, but they do so indefinitely [5]. After establishing the high regenerative capacity of *G.sp*.(Guanajuato), we sought to examine the number, distribution, and functional properties of *G.sp*.(Guanajuato) stem cells.

To assess the functional conservation of *G.sp*.(Guanajuato) stem cells, we turned to a well-established assay of planarian stem cell function. In other highly regenerative planarians, including *S.med*, it is known that neoblasts respond to ionizing radiation in a reproducible and dose-dependent manner [27,28]. Doses ≥ 6000 (6K) rads of γ-radiation cause rapid depletion of *S.med* stem cells, which then leads to homeostasis failure and death. To address whether *G.sp*.(Guanajuato) stem cells respond similarly to □-radiation, we exposed both *G.sp*.(Guanajuato) and *S.med* animals to 10K rads of □-radiation and measured its effects (Figure 6). Notably, most *S.med* animals lysed as expected within 23 days days post irradiation (dpi; Figure 6A,C). However, irradiated *G.sp*.(Guanajuato) maintained normal morphology for nearly 28 dpi and the majority did not lyse until 35 dpi (Figure 6B,C). Importantly, these results were repeated in multiple independent experiments and consistently showed the same animal-specific effect.

After observing such a significant phenotypic difference between *G.sp*.(Guanajuato) and *S.med* in response to lethal □-radiation, we sought to determine the direct effects of this treatment on stem cell number and distribution within *G.sp*.(Guanajuato) animals. To accomplish this, we created a molecular marker of *G.sp*.(Guanajuato) stem cells by cloning the *G.sp*.(Guanajuato) homolog of *piwi-1*, a gene that is highly expressed and enriched in the stem cells of other planarians [25,29]. We generated a fluorescent riboprobe to this sequence and performed *in situ* hybridization in fixed *G.sp*.(Guanajuato) worms. As seen in Figure 6E (0dpi), *G.sp(Gua)_piwi-1* probe not only detects an abundant population of putative stem cells, but the distribution of these cells is very similar to that of *S.med piwi1^+^* cells (Figure 6D, 0dpi), including their absence from the anterior tip of the head and the pharynx.

To analyze the effect of lethal □-radiation on *G.sp(Gua)_piwi1^+^* stem cells, we fixed animals at multiple dpi and performed *in situ* hybridization with both specific *piwi-1* riboprobes (Figure 6E). Intriguingly, lethally irradiated *G.sp*.(Guanajuato) animals maintained a significant number of *G.sp(Gua)_piwi1^+^* stem cells for at least 8 days after lethal □-radiation (Figure 6E, F), whereas *S.med_piwi1^+^* stem cells were rapidly depleted within 2 days as expected (Figure 6D, F). Although this finding did correlate with the relative difference in survival of *G.sp*.(Guanajuato) animals compared to *S.med* (Figure 6A-C), we were still surprised that a small but significant number of *G.sp(Gua)_piwi1^+^* stem cells persisted for so long after such a high dose of □-radiation. We note that the density of *piwi-1^+^* stem cells is too high in the non-irradiated animals (0dpi) for accurate quantitation and analysis as indicated by the “N/A” above this point in the plot.

### Molecular Characterization

Given that the morphological and functional analyses of *G.sp*.(Guanajuato) animals revealed several interesting and distinct features, we proceeded to characterize them molecularly. Our first step was to determine their ploidy using a recently optimized protocol for chromosome preparation and karyotyping in planarians and other organisms [30]. After treating *G.sp*.(Guanajuato) tissue with colchicine to block cycling cells in metaphase, we then swelled, fixed, squashed, and stained the DNA to visualize the metaphase chromosomes. Optimal spreads revealed individual spreads from single cells with clear boundaries and distinct, non-overlapping chromosomes. From these spreads, we identified four diploid chromosome pairs per *G.sp*.(Guanajuato) metaphase cell (Figure 7A). Three previously characterized *Girardia* species are also documented as having 4 chromosomes: *Girardia schubarti, Girardia jenkinsae, and Girardia arizonensis* (Supplemental Table 2). Different from our *Girardia* animals, *Girardia schubarti* is reported to have auricles that are conspicuously nonpigmented [31], which is not the case with our samples (Figure 2A).

Further distinction between *Girardia* species requires either characterization of mature sexual animals or further molecular characterization. However, we do not clearly or reliably observe mature sexual organs in the laboratory colony of this isolate (see Figure 4 and accompanying text). We therefore generated a *G.sp*.(Guanajuato) transcriptome to: 1) validate the *Girardia* genus classification with molecular data; and 2) generate a useful resource for future studies. To create this dataset, we isolated total RNA from a single *G.sp*.(Guanajuato) animal and reverse transcribed an mRNA library using polyA selection. After sequencing and assembling reads from this library, we analyzed the resulting transcriptome for completeness using BUSCO analysis, or “Bench-marking Single Copy Orthologs” [32]; (Figure 7B). The assembled *G.sp*.(Guanajuato) transcriptome is largely complete, as approximately 90% of the 978 metazoan BUSCOs matched completely to assembled *G.sp*.(Guanajuato) transcripts. We then asked how this newly assembled transcriptome compared with those of other planarian species. We downloaded eight additional transcriptomes from the PlanMine database [33] and analyzed each set of assembled transcripts using the same BUSCO parameters as used for our *G.sp*.(Guanajuato) transcriptome analysis. As shown in Figure 7C and Supplemental Table 3, the *G.sp*.(Guanajuato) transcriptome is comparable to that of other planarian transcriptomes, including those that have been experimentally validated in many publications [33–35] in terms of completeness.

After generating this new *G.sp*.(Guanajuato) transcriptome, we compared specific and complete gene orthologs sequences from this isolate with those of multiple flatworm species and created a phylogenetic tree (Figure 7D; Supplemental Figure 5; see also Methods). This analysis clearly supports our classification of the isolate *G.sp*.(Guanajuato) as *Girardia* because it clusters with *Girardia dorotocephala* when analyzed with 10 other flatworm species representing 6 different genera. We also shared the new *G.sp*.(Guanajuato) transcriptome with the Rink lab, who carried out a comparative transcriptome analysis relative to their unpublished transcriptomes. They found significant sequence divergence between the *G.sp*.(Guanajuato) isolate and all other *Girardia* and *Dugesia* isolates in their extensive planarian species collection.

## DISCUSSION

Planarians are best known for their incredible regenerative abilities and negligible senescence, both of which depend on the maintenance of a dynamic and heterogeneous population of adult stem cells. Although some previous studies have identified species-specific differences that underlie regeneration competence in planarians [22–24], comparative studies are surprisingly uncommon in the planarian literature. This is likely due, in large part, to the enduring difficulty of assembling planarian genomes and the resulting lack of robust genomic resources for more than one planarian species. However, although the assembly of planarian genomes has proven difficult in the past [35], recent advances in planarian genome assembly [36] and bioinformatic tools that address genome heterozygosity [37] suggest that this problem is indeed tractable and will pave the way for future genome assembly and comparison projects across multiple planarian species.

Here we characterize a recently collected planarian isolate, *G.sp*.(Guanajuato). Based on both the appearance of all animals at the collection site and years of observation of thousands of animals in the laboratory, most *G.sp*.(Guanajuato) appear to maintain an asexual status, although we cannot fully exclude the possibility that they can be sexualized in specific contexts as seen in other *Girardia* planarians. For example, both *G. tigrina* and *G. dorotocephala* alternate their reproductive status seasonally in nature, becoming sexually mature in cooler months and remaining asexual in warmer months [38]. Both *G. tigrina* and *G. dorotocephala* can also be induced to sexualize in the laboratory upon being fed ground up tissue from the exclusively sexual *Polycelis nigra* [39,40]. However, our attempts to induce sexualization by lowering the temperature or feeding *G.sp*.(Guanajuato) with sexual animal tissue did not induce any observable change in morphology or reproductive function. We did observe a small number of *G.sp*.(Guanajuato) animals that exhibited some evidence of spontaneous sexualization, including the development of ovaries and testes. Spontaneous sexualization of *G. tigrina* laboratory animals has also been reported on multiple occasions [17,41] and these specimens share some similarities with the rare sexualized *G.sp*.(Guanajuato) animals we studied, such as an immature copulatory apparatus and sterility. Nevertheless, the reports of spontaneously sexualized *G. tigrina* describe features that we did not observe in our limited number of putatively sexualized *G.sp*.(Guanajuato), such as hyperplasic ovaries and the presence of supernumerary copulatory apparatuses. Together, our data suggests that *G.sp*.(Guanajuato) may be less capable of sexual maturation than other *Girardia* species or that its induction requires specific, but unknown, environmental inputs.

Analysis of the *G.sp*.(Guanajuato) karyotype also distinguishes this isolate from most other described *Girardia* species (Supplemental Table 2) with two exceptions. Of those species reported to have karyotypes of n=4, those with the greatest similarity to *G.sp*.(Guanajuato) are the North American species *G. arizonensis* [42] and *G.jenkinsae* [43]. Like *G.sp*.(Guanajuato), these species have acutely triangular heads, long auricles, and lightly pigmented pharynges with well-developed inner circular muscle fibers [43]. *G. jenkinsae* and *G. arizonensis* only differ from each other with respect to sexual morphology as determined by examination of histological sections and collection location [43]. Notably, the original report of *G. arizonensis* [42] and the published report of *G. jenkinsae* [43] each mention asexual animals that are morphologically and karyotypically similar to *G. jenkinsae* and *G. arizonensis*, respectively, but are not further characterized in either paper, making it difficult for us to know how they compare to the *G.sp*.(Guanajuato) isolate. We attempted to reconcile whether the *G.sp*.(Guanajuato) isolate and conditionally asexual *G. arizonensis* are distinct species by traveling to the two original collection sites to collect *G. arizonensis* animals. However, forces of both man (fences) and nature (landslide in 1992) made it impossible to access these exact sites, and nearby water sources yielded no specimens in February of 2021.

Although asexual animals are often collected, per taxonomic standards, only sexual specimens are analyzed and described in the reports of new planarian species. The lack of an acceptable standard for classifying distinct asexual species is a major problem with the taxonomic standard. Asexual isolates are either classified as immature animals of a known sexual species (often based on superficial appearance and karyotype) or they are left uncharacterized and underutilized by the research community. There is one notable exception of a named asexual-only species: *Girardia tahitiensis* [44]. Nevertheless, this is only a singular example and the overall underreporting of asexual planarians is detrimental to the advancement of the field, particularly given their demonstrated utility in regeneration research.

One way to overcome the barrier to classifying asexual planarian species is by defining a molecular standard. Although some work has been done comparing the sequences of specific genes, single gene analyses can be misleading and are often insufficient for distinguishing multiple closely related species. As sequencing technology has become increasingly affordable and readily accessible, comparative genomic analyses should become standard in the characterization of new planarian species, as it is the case in many other branches of the tree of life. In fact, there is substantial evidence demonstrating that morphological analyses often underestimate the number of distinct species and can even mislead researchers in characterizing collected specimens [45–48]. Although it is not possible to reanalyze the classification of all published species using transcriptomics, by adding this valuable information to the literature for all current and future species characterizations we can build toward the development of a molecular standard for planarian species classification. This standard would have many advantages, including reducing the variability introduced by individual animals, tissue fixative recipes, and quality of histological sections. It would also allow the field to classify and advance the use of many asexual species that have unique and interesting traits, including *G.sp*.(Guanajuato).

For example, our examination of the *G.sp*.(Guanajuato) revealed two unique, likely interrelated, features: an unusual mode of locomotion and a distinctive pattern of ciliated cells on their outer epithelium. Planarian locomotion has long been known to be propelled by the concerted beating of motile cilia that nearly completely cover the outer epithelium [11,14]. These cilia beat against a layer of secreted mucus, which allows them to achieve a smooth gliding motion. However, *G.sp*.(Guanajuato) appear to have two modes of locomotion: the typical smooth gliding seen in most planarian species and a peristaltic-like motion that is more similar to the “inch-worming” phenotype observed upon chemical or genetic perturbation of motile cilia [18,49]. As this phenotype is a well characterized cilia deficiency, we immediately examined the outer epithelium of *G.sp*.(Guanajuato). Indeed, we found that its appearance is dramatically different from that of other well-studied planarians. The patchy pattern of cilia on the ventral surface of *G.sp*.(Guanajuato) is likely related to its frequent peristaltic-like motion. This pattern is typically observed upon cilia protein depletion (*i.e*., by RNAi), suggesting that *G.sp*.(Guanajuato) may regulate cilia gene expression, cilia protein production, and/or the differentiation of mature epithelial cells differently than other planarian species. The differences in mode of locomotion may also be related to levels of mucus accumulated on the substrate. Given the morphological and functional similarity of this outer mucociliary epithelium to that of many human structures, including the lungs and oviducts, *G.sp*.(Guanajuato) offer an opportunity to identify and study the cellular mechanisms that are dysregulated in various human pathologies affecting these epithelia.

Another way in which *G.sp*.(Guanajuato) distinguish themselves from *S.med* is in their response to high doses of ionizing radiation. We were surprised to observe the significantly long survival time of *G.sp*.(Guanajuato) animals post 10K rads of □-radiation, a dose that is approximately 50 times higher than a standard clinical dose used to treat human cancers. We were even more surprised to observe the persistence of *piwi-1^+^* stem cells for such an extended period of time after radiation treatment, particularly as compared to those of *S.med*. Whereas *G.sp*.(Guanajuato) animals exposed to 10K rads of □-radiation lived approximately 1.5x longer than *S.med* animals treated with this dose, a significant number of *G.sp*.(Guanajuato) *piwi-1^+^* stem cells persisted at least 4x longer than *S.med piwi-1^+^* stem cells. There are many possible explanations for this differential response, including differences in checkpoint mechanisms leading to cell cycle arrest and/or differences in stem cell number. As the cellular response to genotoxic stress has critical implications in human cancer progression and metastasis, studies that dissect the molecular basis of radiation tolerance in *G.sp*.(Guanajuato) are poised to provide important insight into the molecular mechanisms underlying cancer therapy resistance.

The collection and characterization of the planarian isolate, *G.sp*.(Guanajuato), is a valuable addition to the study and documentation of animals in the Turbellaria class. Its unique characteristics raise many interesting questions about their evolution and development, which will undoubtedly spark many future studies and advance our understanding of the cellular and molecular basis of planarian regeneration and stem cell biology

## CONCLUSIONS

Here we describe a newly characterized isolate of planarian flatworm, *G.sp*.(Guanajuato). We show that it is a highly regenerative, putatively asexual, diploid isolate that bears morphological and genetic characteristics of the genus *Girardia*. We document two particular features that suggest it will advance our fundamental understanding of cell biology. By expanding the number and depth of planarian species characterizations, our study will facilitate both comparative studies and those that specifically address the cellular and molecular basis of the unique features of *G.sp*.(Guanajuato).

## METHODS

### Collection and Husbandry

Animals were collected from a site near the Presa San Revato in the Parque Bicentenario in Guanajuato City, Mexico between July 31st and August 2nd of 2016. Animals were transported back to the laboratory and housed in disposable plastic food storage containers filled with Montjuic water [13]. The containers were stored in a dark cabinet at room temperature (22-25C). Worms used in experiments were starved ≥7 days prior to selection for experiments.

### Imaging

All images were adjusted for brightness and contrast using Fiji [50]. High speed imaging of auricle cilia was captured using an ORCA-Flash4.0 V2 C11440-22CU camera from Hamamatsu mounted on a Zeiss Axiovert 200 microscope, as previously described [12]. Light microscopy images of live worm morphology and dimensions were captured with Leica M205 and M165 stereomicroscopes and LAX software. Scaled images were exported and analyzed using Fiji [50].

Scanning Electron Microscopy samples were processed as reported previously [51]. SEM imaging was performed on a Zeiss Merlin. Sectioned samples were processed as in [52], using 0.05 M NaCacodylate buffer. Serial sections were cut on a Leica UC7 ultramicrotome at 100nm section thickness using a Diatome AT-4 35 degree diamond knife. Sections were collected on a glass coverslip and post stained with Sato’s lead for 2 minutes, 4% uranyl acetate in 70% methanol for 3 minutes, and Sato’s lead for 4 minutes. The coverslip was mounted on a stub, coated with 5 nm carbon in a Leica ACE600 coater, and silver paint added to the underside of the coverslip for conductivity. Sections were imaged at 8 kV and 700 pA using the BSD4 detector on a Zeiss Merlin SEM with Atlas 5 software (Fibics, Inc.), and aligned using IMOD [53]. SEM images were processed with the CLAHE plugin (settings: 127, 256, 1.5; available at: https://github.com/axtimwalde/mpicbg/tree/master/mpicbg/src/main/java/mpicbg/ij/clahe). Sectioned EM images used Resize_Replace macro to replace black padding values resulting from image rotation with average grayvale of background (https://github.com/Snowotarski/FUI-macros/tree/20180316_Resize_Replace/Resize_Replace).

### Histology

Tissue processing, section preparation, H&E and special staining were performed as previously conducted in the core lab with modifications [54,55]. Briefly, for tissue processing and section, planarian worms were fixed with 4% PFA in PBS (diluted from 16% (wt/vol) aqueous solution, Electron Microscopy Sciences, cat# 15710) for 24hr at 4°C, then rinsed well with 1xPBS, dehydrated through graded ethanol (30%, 50%, 70%) and processed with a PATHOS Delta hybrid tissue processor (Milestone Medical Technologies, Inc, MI). Using a Leica RM2255 microtome (Leica Biosystems Inc. Buffalo Grove, IL), sections were cut in 5 μm thickness and mounted on Superfrost positive charged microscope slides (cat# 12-550-15, Thermo Fisher Scientific). H&E routine stain was performed using Leica ST5010 Autostainer XL and Leica ST Infinity H&E Staining System kit (VWR#3801698) according to the manufacturer’s instructions. Resultant slides were imaged on an Olympus America Slide Scanner.

### In-situ Hybridization

In-situ hybridization was performed as previously described using Digoxigenin labeled riboprobes and tyramide amplification of their signal [56,57]. After probe hybridization, signal development, and mounting on slides, animals were imaged on a CSU-W1 with a Nikon base and Prior plate loading robot as described [58]. Fully automated image acquisition was obtained by first imaging whole slides using a 4X 0.2NA objective. Individual worms were segmented and z-stacks acquired using a 10X 0.45NA objective. Stitching and spot quantification were performed in Fiji using custom macros (https://github.com/jouyun/smc-macros).

### Irradiation

Irradiation was done with a Gammacell 40 Exactor (MDS Nordion) at a dose rate of 85 Rad/minute.

### Karyotyping

Karyotyping was carried out as previously described [30].

### RNA Extraction and Sequencing

Total RNA from one worm was extracted with TRIzol™ (Invitrogen) using the standard protocol. An mRNAseq library was generated from 100ng of high quality total RNA, as assessed using the Bioanalyzer (Agilent). The library was made according to the manufacturer’s directions for the TruSeq Stranded mRNA LT Sample Prep Kit - set B (Illumina, Cat. No. RS-122-2102). The resulting short fragment library was checked for quality and quantity using the LabChip GX (PerkinElmer) and Qubit Fluorometer (Life Technologies). The library was then sequenced as 100bp paired reads on two rapid-run flow cells using the Illumina HiSeq 2500 instrument. Following sequencing, Illumina Primary Analysis version RTA 1.18.64 and bcl2fastq2 v2.18 were run to demultiplex the sequencing reads and generate FASTQ files.

### Transcriptome Assembly

Transcriptome assembly was created with Trinity (version: v2.2.0; parameters: --trimmomatic -- min_contig_length 300) [59]. Post-assembly vector trimming and contaminate filtering was performed with SeqClean (version: 20110222; parameters: -l 300) [60]. Transcriptome assembly consists of 183,422 sequences with a mean GC content of 32%.

Sequences were translated with Transdecoder (version: v2.0.1) [61] and collapsed into longest isoforms with cd-hit (version: 4.6, parameters: -c .95 -G 0 -aS .5) [62]. The resulting non-redundant file of predicted proteins consisted of 23,850 proteins of which 10,303 contain both start codons and stop codons; 5,993 contain stop codons, but not start codon; 2,409 contain a start codon, but not a stop codon; and 5,145 contain neither stop, not start codons.

### Transcriptome Completeness Assessment

BUSCO (version: 3.1.0; parameters: -l metazoa_odb9) [32] was used for completeness analysis. BUSCO found 89.5% of metazoan single-copy orthologs in the *G.gua* transcriptome assembly with only 8.8% of single-copy orthologs missing (summary: C:89.5%[S:60.5%,D:29.0%],F:1.7%,M:8.8%, n:978). BUSCO analyses of other planarian transcriptomes were performed on transcriptomes downloaded from Planmine (Supplemental Table 3).

### Phylogenetic Analysis

Preliminary BLAST [63] analysis suggested *G.sp*.(Guanajuato) as most closely related to a previously published *Girardia* species. To refine the position of this species in the planarian phylogenetic tree OrthoMCL (version: v2.0.9) [64] was used to define putative ortholog groups from *G.sp*.(Guanajuato) and 11 published transcriptomes: *Dendrocoelum lacteum, Polycelis nigra, Polycelis tenuis, Schmidtea mediterreanea, Planaria torva [33], Dugesia japonica* (GFJY01000000.1), *Schmidtea polychroa* (GFKA00000000.1), *Girardia dorotocephala* (GEIG01000000.1), *Phagocata morgani* (GEKK01000000.1), *Phagocata gracilis* (GEGP01000000.1) and *Bdelloura candida* (GFKB01000000.1).

Gene groups with no more than one sequence from each species and present in at least ten of the selected species were separately aligned with MUSCLE (version: v3.8.31) [65], trimmed with trimAl(version: v1.4.rev22; parameters: -automated1) [66] and then concatenated and cleaned with phyutility (version: v.2.2.6; parameters -clean .5) [67]. Phylogenetic tree was created using RAxML-NG (version: 0.8.1 BETA; parameters: --model JTT+G+F --tree pars{10} --seed 123 --bs-trees 200) [68].

## List of abbreviations

*G.sp*.(Guanajuato): *Girardia species isolate from Guanajuato*
S.med: *Schmidtea mediterranea*
Dpa: days post amputation
Dpi: days post irradiation

---

**Supplemental Figure 1.**
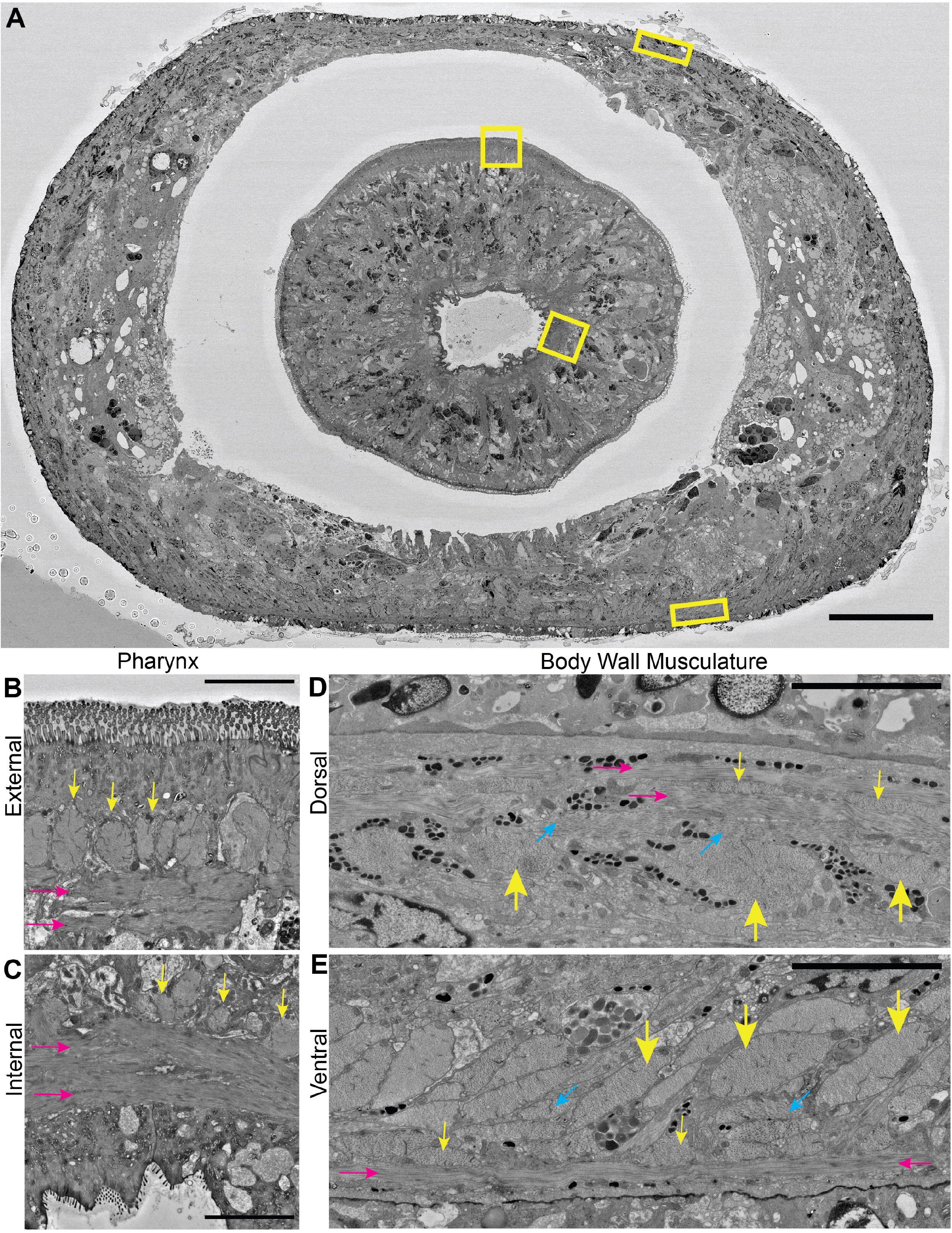
Muscle layers in *G.sp*.(Guanajuato) (A)Low magnification SEM image of a transverse cross section of *G.sp*.(Guanajuato) through the middle of the pharynx.(B) The external muscle layer of the pharynx is a layer of longitudinal muscles (yellow arrows) immediately superficial to a deeper circular muscle layer (magenta arrows). (C) The internal muscle layer in the pharynx has a deep layer of longitudinal muscles, and a more superficial and thicker layer of circular muscle fibers. (D)The body wall musculature is a thin circular muscle layer (magenta arrows) with a deeper thin longitudinal fiber layer (thin yellow arrows), under which is a diagonal muscle layer (cyan arrow) which is underlain by a much thicker layer of longitudinal fibers (thick yellow arrows)(E) The body wall musculature is somewhat more developed on the ventral side, especially the inner longitudinal muscle fibers. Scale Bars = 100μm in A, 5μm in B-E.

**Supplemental Figure 2.**
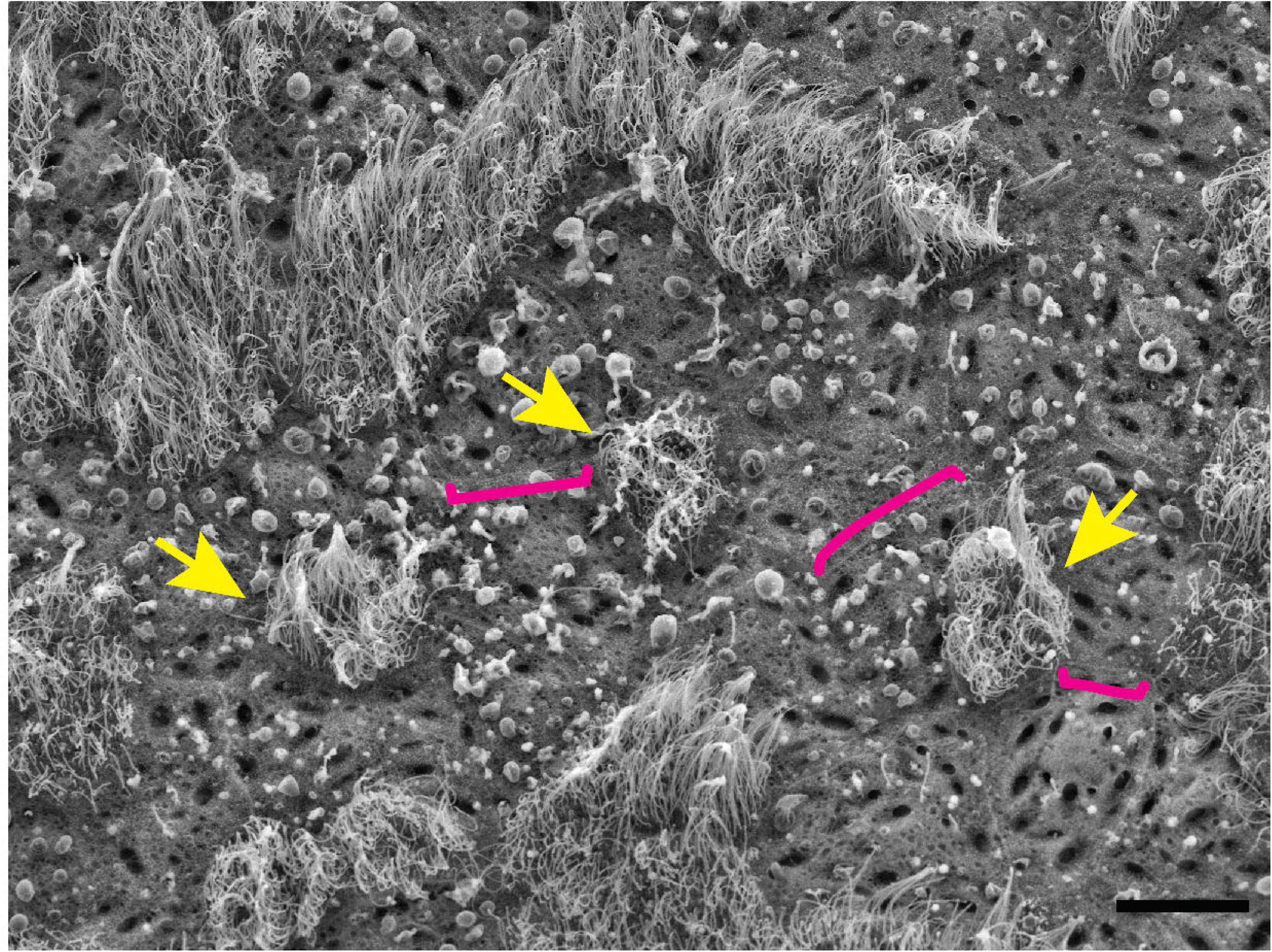
Single ciliated cells on the ventral epidermis of *G.sp*.(Guanajuato) High magnification SEM of field of ventral epidermis cells. Patches of ciliated cells are evident in the top left while single ciliated cells in the ventral epidermis (yellow arrows) are midfield. Cell-cell junctions (magenta brackets) mark cell boundaries. Scale bar = 10μm

**Supplemental Figure 3.**
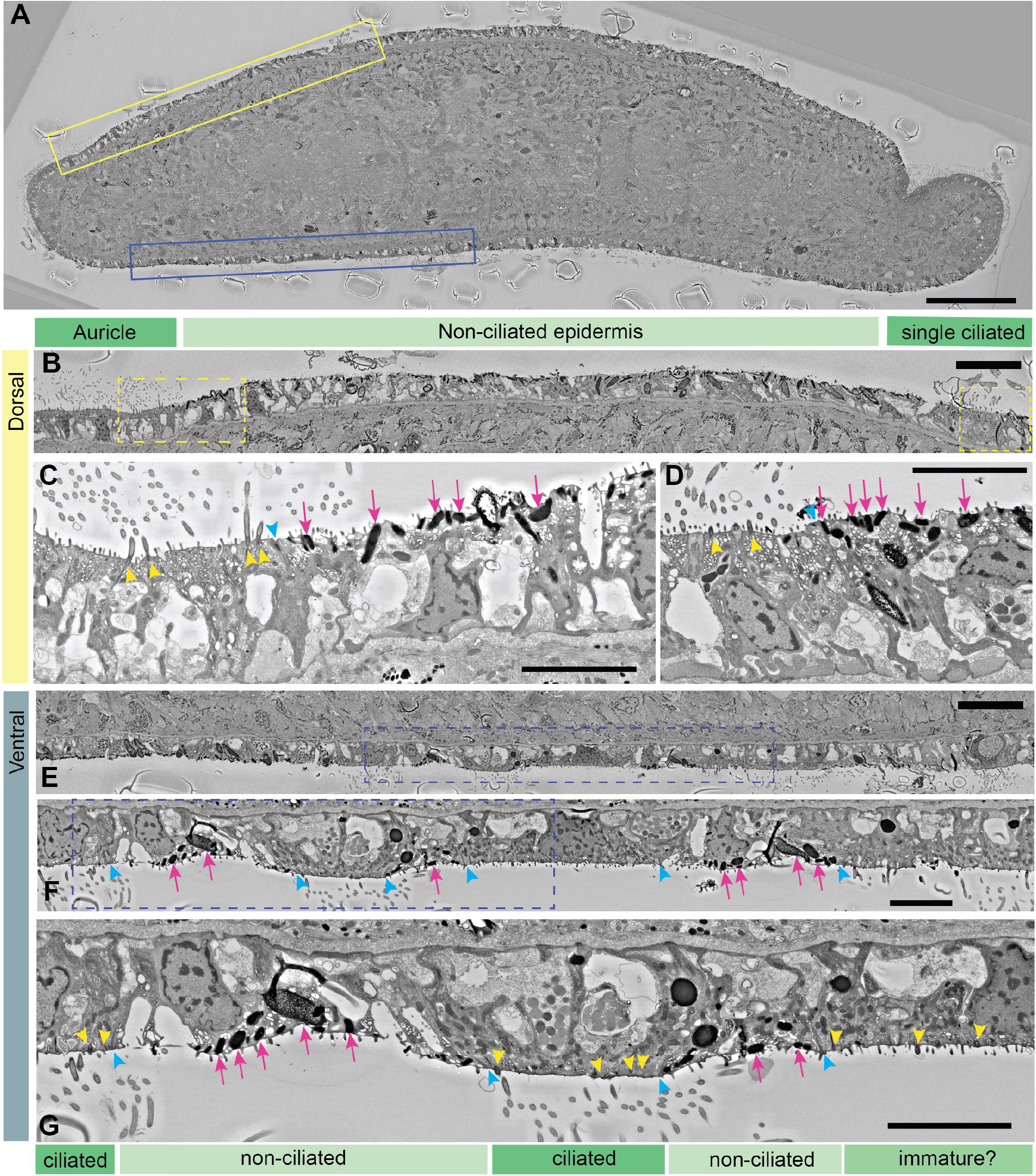
Differences in ciliated and non-ciliated epithelial cells in *G.sp*.(Guanajuato) (A)Low magnification Scanning Electron Microscopy (SEM) image of a transverse cross section of *G.sp*.(Guanajuato) through both auricles with dorsal and ventral regions further depicted in later panels outlined.(B) As expected from the SEM images, the dorsal epidermis is largely non-ciliated with the exception of ciliated cells at both the lateral auricles and at medial single cells (ciliated regions marked in dark green bars, non-ciliated regions in light green). (C)Closer inspection of the auricles reveals basal bodies associated with cilia (yellow arrowheads), while just across the cellcell junction at the auricle boundary(cyan arrowhead), the non-ciliated epidermal cells do not have basal bodies, but do however clearly contain many rhabdites (magenta arrows). (D)The single ciliated cells close to the midline also have the same features: cilia, basal bodies and lack of rhabdites. (E) The ventral epidermis, as anticipated from SEM images, contain ciliated and non-ciliated cells. (F) Using cell-cell junctions (cyan arrowheads) as boundaries it is clear that ciliated cells do not contain rhabdites, while non-ciliated cells do, as in the doral epithelium.(G) Ciliated cells contain basal bodies(yellow arrowheads), while the majority of non-ciliated cells do not. Few instances of cells containing basal bodies but not full cilia are likely immature ciliated cells. Scale bars = 100μm in A, 25μm in B and E, and 5μm in C,D,F and G.

**Supplemental Figure 4.**
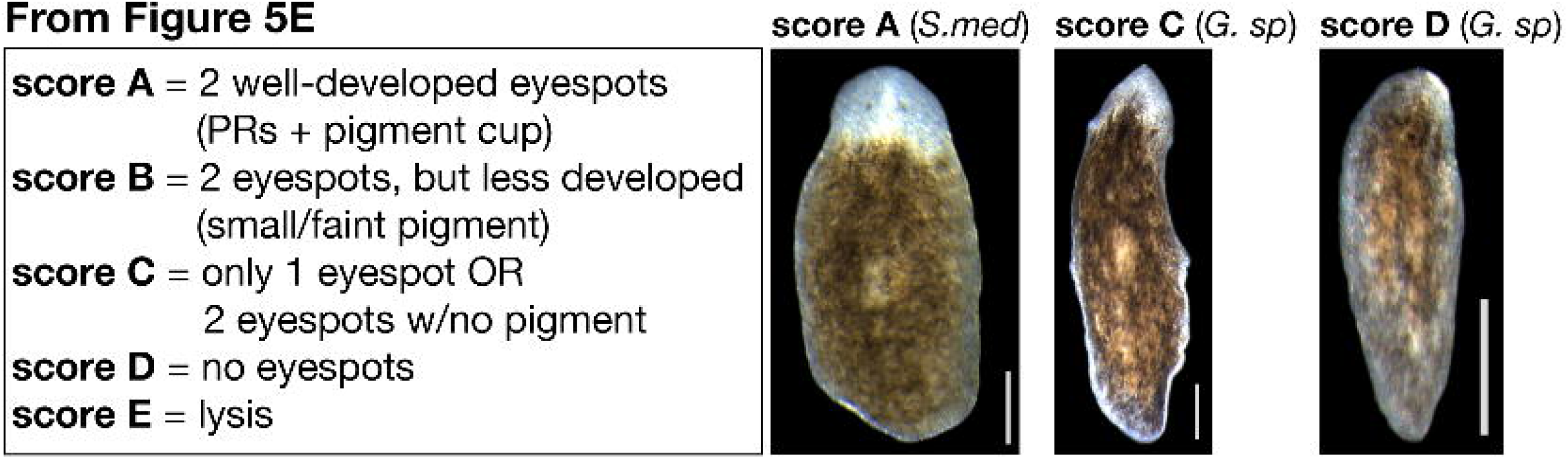
*G.sp*.(Guanajuato) regenerate differently from *S.med*. Representative images of regenerating *S.med* and *G.sp*.(Guanajuato) fragments from the experiment summarized in Figure 5D-E.

**Supplemental Figure 5.**
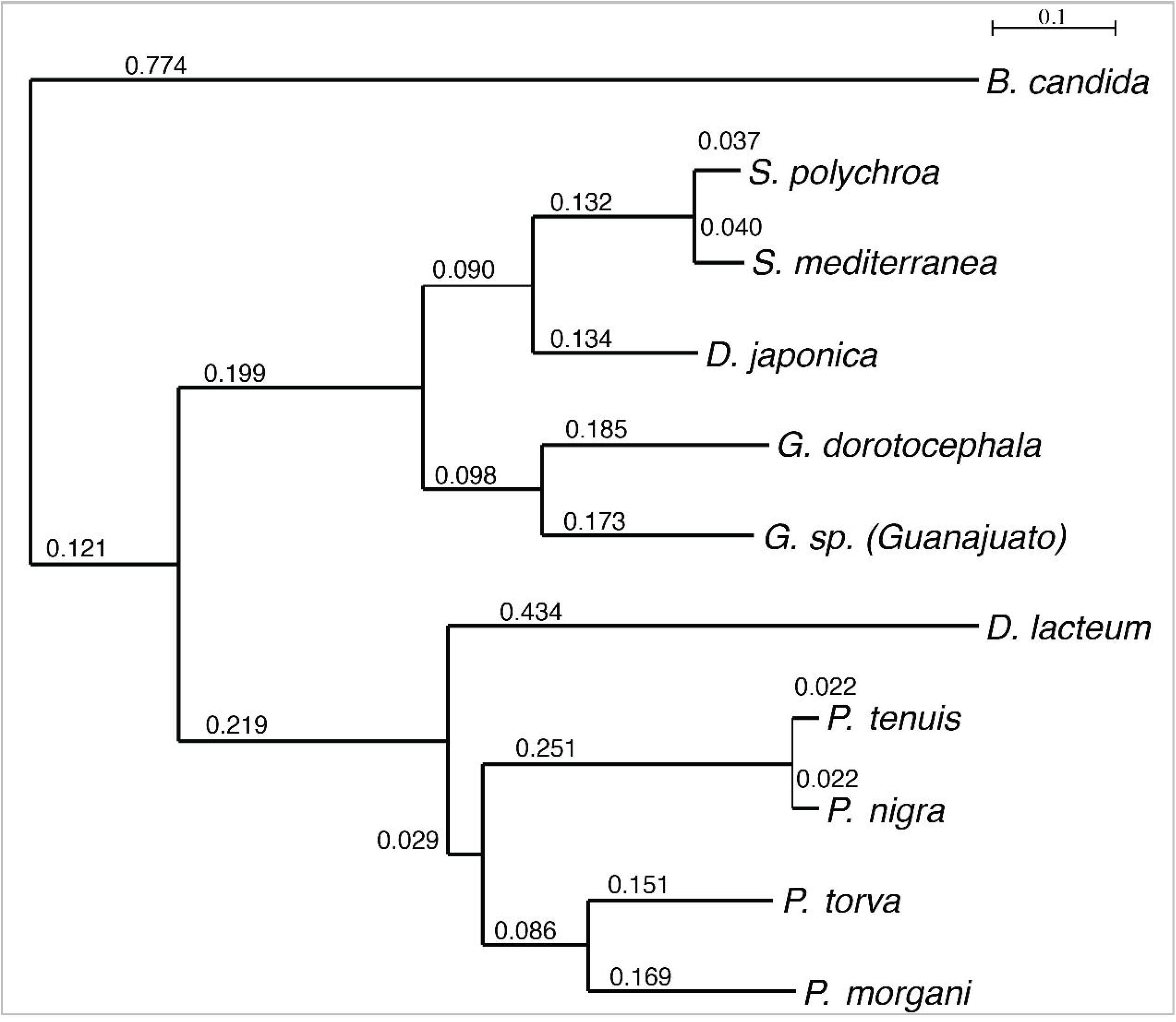
*G.sp*.(Guanajuato) are a *Girardia* species. Phylogenetic tree from Figure 7D with scale bar units labeled for branch lengths. Scale bar unit = expected number of substitutions per site.

## DECLARATIONS

### Ethics approval and consent to participate

Not applicable.

### Consent for publication

Not applicable.

### Availability of data and materials

Original data underlying this manuscript can be accessed from the Stowers Original Data Repository at http://www.stowers.org/research/publications/libpb-1535. The *G.sp*.(Guanajuato)transcriptome is deposited with the NCBI Bioproject: PRJNA551297. NCBI assigned *G.sp*. (Guanajuato)with taxonomy ID: 2592332 and name Girardia sp. N. ER-2019.

### Authors’ contributions

ASA and CdeC located and collected the specimens. EMD, SHN, CGH, and EJR contributed to experimental design. CGH completed histological specimen preparations, and RNA isolation. JD completed the regeneration experiment and live imaging of regenerating fragments. CGH and EMD completed live worm imaging, irradiation, and stem cell staining experiments. SM performed all fluorescent imaging and image analyses. SHN performed SEM and pharynx imaging. LG and CGH performed all chromosome preparations and karyotyping. EJR completed phylogeny and transcriptome analyses. EMD and SHN wrote the manuscript.

## Acknowledgements

We thank all members of the Sánchez Alvarado Lab for discussion and insightful comments. We thank the Histology Core and the Light and Electron Microscopy Core at Stowers Institute for Medical Research for SEM sample preparation, histological preparation, and slide scanner imaging. We particularly thank Yongfu Wang for assistance with histology and Jeff Lange for his assistance in capturing the live, high speed images of beating cilia. We also acknowledge the assistance of Sofia Robb at the Stowers Institute for Medical Research for help with OrthoMCL, as well as Brian Fleharty and the Stowers Institute for Medical Research Molecular Biology Core facility for library preparation and sequencing. We also extend thanks to Marta Riutort for integral comments regarding Girardia.

In 2016, the Clubes de Ciencia México Workshop for Developmental Biology included: Leslie Yareli Govea-Martínez, Cristell Hernández-Fonseca, Benjamín Pérez-Sánchez, Israel Torres-Alba, Maria Ramirez-Morales, Erick Navarro-Delgado, Nohemí Carolina García-Rodríguez, Arizbeth Plascencia, José Luis Rodríguez-Ortiz, L. Gerardo Moreno-Ciénega, Adrian Maciel-Avalos, Porfirio Gallegos-Casillas, Mónica Vázquez Guerrero, Eduardo Zamudio de la Cruz, Marcos Romero-Partida, Rodrigo Jonathan Martinez-Espinosa, Abigail Zuñiga-Arenas, Sofía Quiroz Yebra, Sara Elena Ambriz-Piña, José Segoviano, Karla Fernández (undergraduate students); Fernando Longoria Vázquez (course assistant); Heather Leigh Curtis and David Angeles-Abores (course organizers).

## Additional Files

⍰ supplemental movie S1 (movie, .mp4); wild-type *G.sp*.(Guanajuato)worm motility

⍰ supplemental movie S2 (movie, .mp4); motile cilia on auricle of wild-type *G.sp*.(Guanajuato)worm

⍰ supplemental movie S3 (movie, .mp4); serial sections of copulatory region

⍰ supplemental figure 1 (.png); SEM image of section detailing the pharyngeal and body wall muscle

⍰ supplemental figure 2 (.png); high magnification SEM image showing single ciliated and non ciliated cells

⍰ supplemental figure 3: (.png); SEM image of cross section through the auricular region detailing features of the outer *G.sp*.(Guanajuato) epithelium

⍰ supplemental figure 4: (.pdf); Phylogenetic tree as in Figure 7D labeled with node values

⍰ supplemental table 1: (.png); body length and pharynx measurements for 8 animals

⍰ supplemental table 2: (.xlsx); compilation of data on *Girardia* species collected in Mexico or with a karyotype of n=4

⍰ supplemental table 3 (BUSCOs) (pdf, .pdf); BUSCO Results for all transcriptomes in Figure 7C

