## Supplementary figures and images for "Molecular characterization of a flatworm Girardia isolate from Guanajuato, Mexico"

### Supplemental Table 1

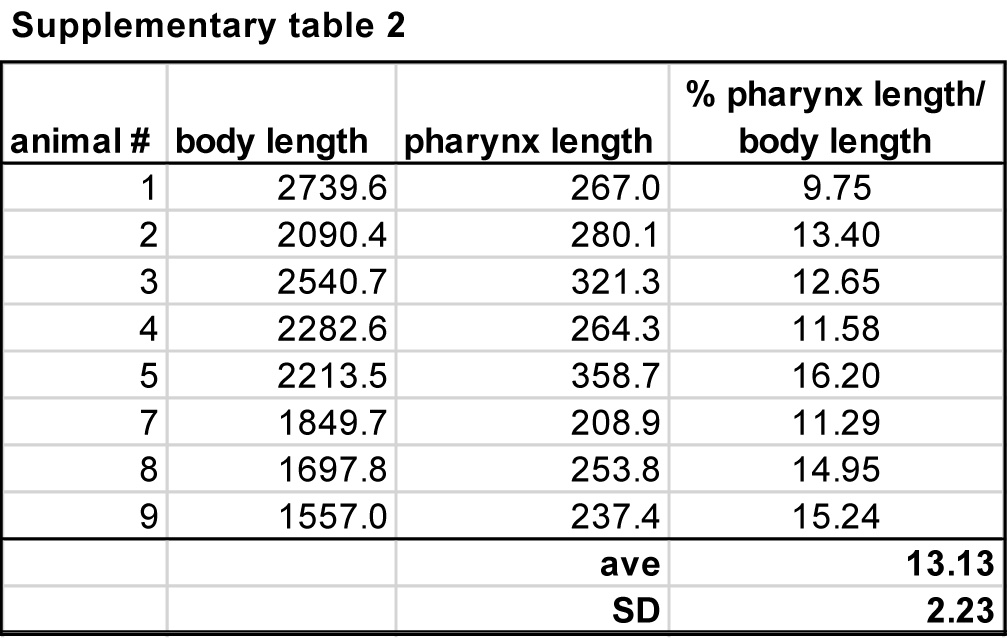
